# Control of assembly of extra-axonemal structures: the paraflagellar rod of trypanosomes

**DOI:** 10.1101/2019.12.17.879841

**Authors:** Aline A. Alves, Maria J. R. Bezerra, Wanderley de Souza, Sue Vaughan, Narcisa L. Cunha-e-Silva, Jack D. Sunter

## Abstract

Eukaryotic flagella are complex microtubule based organelles and in many organisms there are extra-axonemal structures present, including the outer dense fibres of mammalian sperm and the paraflagellar rod (PFR) of trypanosomes. Flagellum assembly is a complex process occurring across three main compartments, the cytoplasm, the transition fibre-transition zone, and the flagellum. It begins with translation of protein components, followed by their sorting and trafficking into the flagellum, transport to the assembly site and then incorporation. Flagella are formed from over 500 proteins; the principles governing axonemal component assembly are relatively clear. However, the coordination and sites of extra-axonemal structure assembly processes are less clear.

We have discovered two cytoplasmic proteins in *T. brucei* that are required for PFR formation, PFR assembly factors 1 and 2. Deletion of either PFR-AF1 or PFR-AF2 dramatically disrupted PFR formation and caused a reduction in the amount of major PFR proteins. The presence of cytoplasmic factors required for PFR formation aligns with the concept of processes occurring across multiple compartments to facilitate axoneme assembly and this is likely a common theme for extra-axonemal structure assembly.

**Summary statement:** Eukaryotic flagella are complex organelles. In many organisms there are extra-axonemal structures including the trypanosome paraflagellar rod. We have discovered two cytoplasmic proteins that are required for paraflagellar rod formation.

## Introduction

The eukaryotic flagellum is a well conserved organelle which has multiple functions that include providing a propulsive force and acting as a sensory platform (Moran et al., 2014). A flagellum consists of a microtubule axoneme surrounded by plasma membrane. The number and arrangement of microtubules can vary in the axoneme with a motile flagellum typically containing 9 outer microtubule doublets encircling a central pair of singlet microtubules to give a 9+2 arrangement, whereas a primary cilium (used interchangeably with flagellum) has a 9+0 axoneme, lacking the central pair. Flagellar proteomes of diverse organisms and the recent genome-wide protein localisation project (TrypTag) in *T. brucei* have shown that the flagellum is a complex organelle containing over a 1000 proteins (Beneke et al., 2019; Broadhead et al., 2006; Dean et al., 2017; Hart et al., 2009; Narita et al., 2012; Ostrowski et al., 2002; Pazour et al., 2005).

Flagellum assembly is a complex, multi-site process involving three main compartments, 1) the cytoplasm, 2) the transition fibre-transition zone, and 3) the flagellum. Protein components are synthesised in the cytoplasm, then sorted and directed to the flagellum via the transition fibre and transition zone with many transported to the tip by the intraflagellar transport system before being incorporated into the flagellum structure (Langousis and Hill, 2014; Sánchez and Dynlacht, 2016). Not all flagellar proteins are transported into the flagellum as individual proteins. Dynein arms are large, highly structured complexes (11-17 proteins) comprising heavy, intermediate and light chains that are attached to the A tubule of the outer microtubule doublet of the axoneme (Kobayashi and Takeda, 2012). Mutations in dynein proteins have been shown to cause defective swimming in *Chlamydomonas* and primary ciliary dyskinesia in humans (Desai et al., 2018; Kobayashi and Takeda, 2012). In addition, to the dynein proteins themselves disruption of other proteins can cause the loss of the axonemal outer and inner dynein arms, resulting in flagellar motility defects. Investigation of these proteins predominantly in *Chlamydomonas* has led to the discovery of an ordered axonemal dynein assembly process that has three key steps each occurring in each of the three main compartments: 1) cytoplasmic assembly and maturation of the outer dynein arm complex, 2) transport of the complex into the flagellum, and 3) docking of the complex to the microtubule doublet (Desai et al., 2018). Thus whilst some components of the axoneme seem to travel to the flagella tip as individual proteins, others are pre-assembled in the cytoplasm; however, all are influenced by the three main compartments required for flagellum assembly.

In many organisms there are additional extra-axonemal structures; for example, the outer dense fibres and fibrous sheath in mammalian sperm, mastigonemes in *Chlamydomonas*, vane structures found in protists such as the fornicate *Aduncisulcus paluster* and the paraflagellar rod (PFR) in *Trypanosoma brucei* and other *Euglenozoa* (de Souza and Souto-Padrón, 1980; Hyams, 1982; Irons and Clermont, 1982a; Irons and Clermont, 1982b; Nakamura et al., 1996; Portman and Gull, 2010; Yubuki et al., 2016). When viewed by thin section electron microscopy the outer dense fibres, the ventral vane of *A. paluster* and the PFR all have a striated appearance, suggesting a regular high-order structure (Farina et al., 1986; Portman and Gull, 2010; Woolley, 1971; Yubuki et al., 2016). The sperm outer dense fibres are associated with the 9 outer microtubule doublets in the principal piece of the sperm flagellum (Irons and Clermont, 1982a). Surrounding the axoneme and the outer dense fibres is the fibrous sheath that is formed of two longitudinal columns, which are attached to outer dense fibres 3 and 8 and are connected to each other by semi-circular transverse ribs (Eddy et al., 2003). The outer dense fibres contain at least 25 proteins with a further 9 known to localise to the fibrous sheath (Eddy et al., 2003; Petersen et al., 1999). The predominate proteins in the fibrous sheath are two A-kinase anchoring family proteins (AKAPs), AKAP3 and AKAP4, which enable the fibrous sheath to act as a platform for signalling and metabolic pathways. Associated with the fibrous sheath are many proteins involved in glycolysis e.g. isoforms of GAPDH and HK1 (Eddy et al., 2003). Disruption of the expression of outer dense fibre proteins such as ODF2 and fibrous sheath proteins including AKAP4 caused defects in outer dense fibre and fibrous sheath structure that impacts sperm motility (Miki et al., 2002; Tarnasky et al., 2010; Zhao et al., 2018). These structures therefore likely provide mechanical support and also act as signalling and metabolic platforms, important for flagellar beat regulation.

Both the outer dense fibres and the fibrous sheath are assembled in the sperm flagellum once the axoneme has been built. The outer dense fibres are built in a proximal to distal direction along the flagellum, whereas the fibrous sheath is assembled in a distal to proximal direction with the longitudinal columns assembled first before being connected by the transverse ribs (Irons and Clermont, 1982a; Irons and Clermont, 1982b). Little is known about the mechanism of the assembly of these structures; however, the deletion of ubiquitin conjugating enzyme UBE2B resulted in sperm flagella that had a normal axoneme structure but disrupted positioning of the longitudinal columns (Escalier, 2003). In *Chlamydomonas* during flagella regeneration, mastigonemes appeared on the new flagellum within 15 minutes of amputation of the old flagella, suggesting that these structures as with the outer dense fibres and fibrous sheath are assembled after the axoneme. Moreover, the mastigonemes were not found at the base of the newly assembled flagellum but instead on the distal three quarters of the regenerating flagellum (Nakamura et al., 1996).

In *Euglenozoa*, including *T. brucei*, the axoneme is accompanied by the extra-axonemal PFR. In *T. brucei* nearly 200 proteins have been found in the PFR, including the two most abundant components, PFR1 and PFR2 (Dean et al., 2017; Portman et al., 2009). The PFR contains proteins such as adenylate kinases, cyclic nucleotide phosphodiesterases, and calmodulin indicating it has a role in both cAMP and calcium regulation that are likely to be relevant to flagellum beat regulation and to possible sensory functions, hence showing parallels to the role of the fibrous sheath in sperm flagella (Ginger et al., 2013; Luginbuehl et al., 2010; Moran et al., 2014; Pullen et al., 2004).

Normally, the PFR lies parallel to the axoneme with the structure first appearing at a variable distance from the basal body depending on the species and then tapering towards the flagellum tip. The PFR has an intricate paracrystalline structure with three distinct domains (proximal, intermediate, distal) and is attached to the axoneme via microtubule doublets 4 and 7 (Farina et al., 1986; Portman and Gull, 2010). However, the PFR can vary dramatically in structure with *Angomonas deanei* and *Strigomonas culicis* having a very short and simplified PFR (Gadelha et al., 2005; Motta et al., 2013). The extra-axonemal PFR of *T. brucei* is assembled via PFR1 and PFR2 subunit incorporation at the distal tip of the growing flagellum and lags behind axoneme assembly (Bastin *et al*., 1999b). The two most abundant proteins in the PFR, PFR1 and PFR2, are critical for its assembly. Loss of PFR1 and PFR2 components in *T. brucei* or the related kinetoplastid, *Leishmania mexicana* results in a loss of the PFR structure (Santrich *et al*., 1997; Bastin *et al*., 1998; Maga *et al*., 1999; Bastin *et al*., 1999b), and a profound reduction in motility. To date only one other PFR protein, calmodulin has been shown to have an important role in PFR assembly with many minor protein components such as PFC3, PAR1 not required for its assembly (Ginger et al., 2013; Lacomble et al., 2009).

The incorporation of axonemal proteins into the flagellum requires processes within three different compartments (cytoplasm, transition fibre-transition zone, flagellum). We know that extra-axonemal structure assembly occurs within the flagellum of cells such as trypanosomes and sperm (Bastin et al., 1999a; Irons and Clermont, 1982a; Irons and Clermont, 1982b). In trypanosomes two non-PFR proteins have been identified to be important for PFR assembly, Kif9b and FOPL (Demonchy et al., 2009; Harmer et al., 2018). Kif9b is found in the basal body, pro-basal body, and axoneme and FOPL is found at the transition fibres. RNAi mediated knockdown of either of these two proteins caused severe defects in PFR assembly with flagellum sections showing either no PFR or accumulation of multiple PFR units. The position of FOPL at the transition fibres shows that this compartment is also important for PFR formation. Given that PFR formation requires processes within the flagellum and at the transition fibres - are there processes in the cytoplasm required for extra-axonemal formation?

Here, we identified two cytoplasmic proteins, which together form a complex that is required for PFR formation. This suggests that assembly of extra-axonemal structures as with the axoneme also require processes in all three compartments. Presumably, such cytoplasmic localised phenomena reflect the need for construction processes of major structures to be regulated and coordinated to ensure delivery of correct stoichiometric amounts of components at precise temporal windows and via particular transport systems.

## Results

### *Tb927.10.8870* deletion causes slow growth and errors in cytokinesis

Given the complex structure of the PFR, we hypothesised that additional proteins would be required for its formation. We investigated Tb927.10.8870, a ∼34 kDa protein, which is predicted to be a coiled coil protein that has identity to the taxilin PFAM domain. Results from TrypTag, a global genome survey of protein localisation showed that the mNeonGreen (mNG) tagged Tb927.10.8870 protein localised to the cytoplasm and was excluded from the flagellum and the nucleus (Dean et al., 2017). We repeated this by endogenously tagging Tb927.10.8870 with mNG at its C-terminus and examining its localisation (Figure 1A). Tb927.10.8870::mNG was restricted to the cytoplasm and had a patchy distribution as seen previously, which was concentrated in the posterior half of the cell.

**Figure 1:**
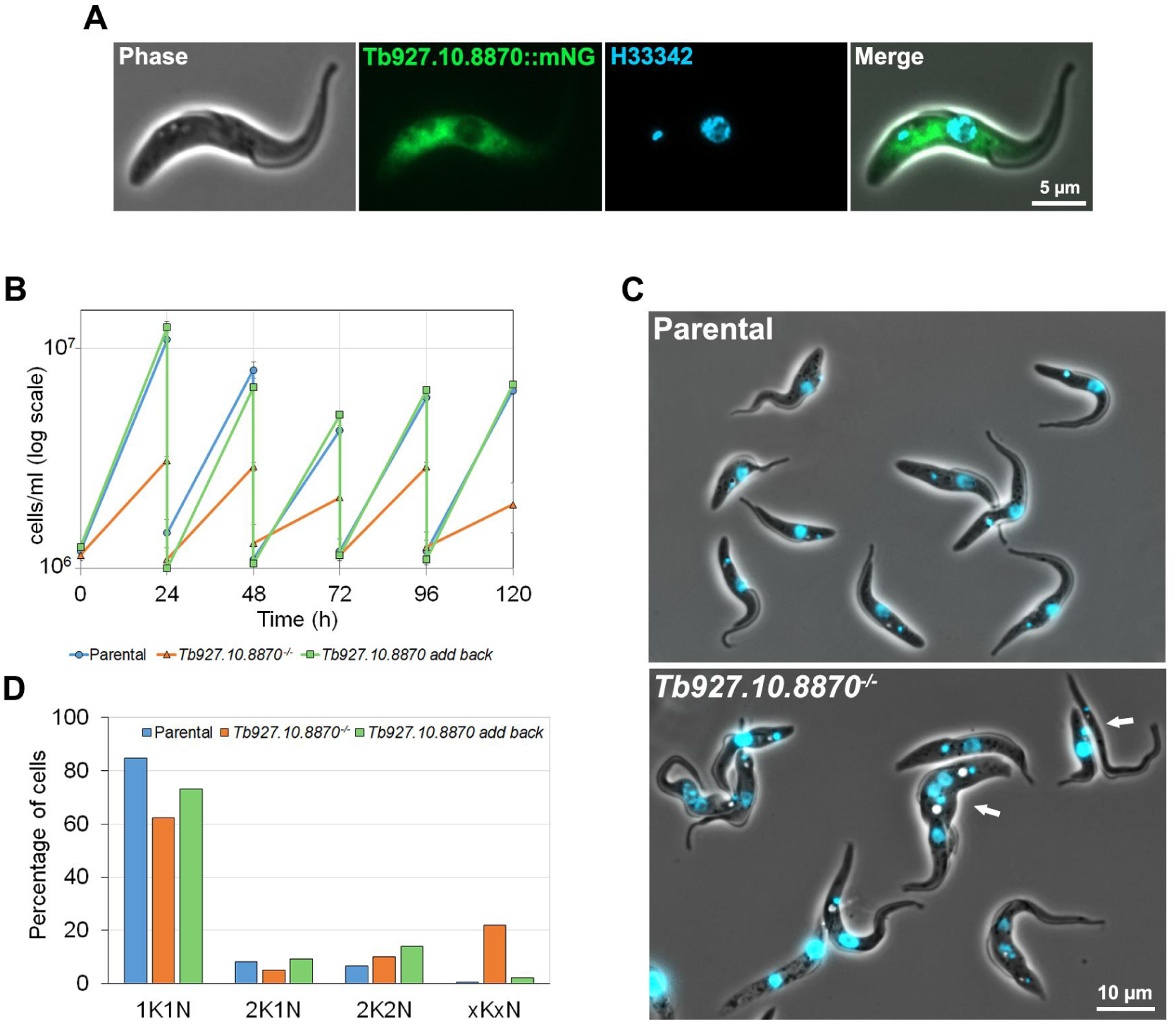
Tb927.10.8870 is a cytoplasmic protein that was required for robust cell growth. (A) Live cell expressing Tb927.10.8870::mNG (green) stained with DNA stain Hoechst 33342 (blue). Scale bar: 5 µm. (B) Growth curves of parental, *Tb927.10.8870*^*-/-*^ and *Tb927.10.8870* add back cell lines over 120 hours. (C) Merge of phase and Hoechst images of parental and *Tb927.10.8870*^*-/-*^ live cells. White arrows indicate abnormal cell types. Scale bar: 10 µm. (D) Nuclear (N) and kinetoplast (K) DNA was counted in 200 cells using Hoechst 33342 stained fluorescence images of parental, *Tb927.10.8870*^*-/-*^ and Tb927.10.8870 add back cell lines. All cells with abnormal numbers of K and N were counted as “xKxN”.

We investigated the function of Tb927.10.8870 by generating a cell line in which both alleles of the gene had been replaced with antibiotic resistance markers using a CRISPR/Cas9 based approach (Beneke et al., 2017). We did this in a newly developed cell line, which expresses the Cas9 nuclease, T7 RNA polymerase and tet repressor from a single plasmid called pJ1339 and is therefore competent for both CRISPR/Cas9 genome editing and tetracycline controlled inducible expression.

We were able to readily generate cell lines that were resistant to both selective drugs and confirmed the loss of both alleles of *Tb927.10.8870* and integration of the markers by PCR (Figure S1A). However, these null mutants took a longer time than normal to grow through after the transfection and we therefore measured the growth rate of the null mutant (Figure 1B). The deletion of Tb927.10.8870 caused the cells to grow consistently slower than the parental cell line. Moreover, by light microscopy a number of abnormal forms were observed (Figure 1C). The reduced growth rate and abnormal cell forms suggested that cytokinesis was potentially disrupted in this cell line. During the trypanosome cell cycle the kinetoplast (concatenated mitochondrial DNA) and the nucleus replicate and divide at specific time points and their relative number in the cell indicates the cell cycle stage. To investigate if the cell cycle was disrupted we imaged the null mutant and parental cells with a fluorescent DNA stain (Figure 1D). There was a reduction in the number of cells with 1 kinetoplast (K) and 1 nucleus (N) in the null mutant and an increase in the number of cells with an abnormal KN number, indicating that there was a defect in cytokinesis in the null mutant.

To confirm that the changes we observed in the null mutant were due to the loss of Tb927.10.8870 we generated a Tb927.10.8870 add back cell line; a copy of the *Tb927.10.8870* gene was introduced into the null mutant with a 5’ Ty epitope tag (Bastin et al., 1996). We confirmed the expression and localisation of Tb927.10.8870 in the add back cell by western blot and fluorescence microscopy (Figure S1B, C). The Ty tagged protein ran at the expected size on the western and was restricted to the cytoplasm as observed with the mNG tagged protein. The growth rate of the add back cell line was similar to that of the parental cell and cells with abnormal KN numbers were much less frequent than compared to the null mutant (Figure 1C, 1D). This shows that the slow growth and cytokinesis defect were due to the loss of Tb927.10.8870 and unlikely due to an off-target effect.

### Tb927.10.8870 is required for PFR assembly

Careful examination of the light micrographs of the null mutant cells showed that 52% (n=300) of cells with one flagellum had a bulge at the distal tip of the flagellum in comparison to only ∼3% for the parental cells (n=92) and the add back cells (n=165) (Figure 2B). When the cells were examined by scanning electron microscopy, the bulge at the tip of the flagellum was readily apparent (Figure S2A). This suggests that flagellum assembly in these cells was disrupted. To determine whether there were changes in the trypanosome flagellum on deletion of Tb927.10.8870 we stained the parental, null mutant and add back cells with monoclonal antibodies to an axonemal component TbSAXO (mAb25) and the PFR, component PFR2 (L8C4) (Figure 2A). In the parental, null mutant and, add back cells mAb25 stained a linear structure in the flagellum that extended from close to the kinetoplast to the tip of the flagellum, correlating with the position of the flagellum axoneme. In the parental and add back cells L8C4 stained a linear structure within the flagellum from the point at which the flagellum exited the flagellar pocket and along the majority of the flagellum before fading towards the distal tip, which corresponds to the location of the PFR. However, in the null mutant the L8C4 signal in the flagellum was no longer evenly distributed with patches of strong staining interspersed with regions of very weak staining and often there was a strong signal at the distal tip of the flagellum that coincided with a bulge. This suggests that Tb927.10.8870 is important for PFR assembly but not axoneme assembly and hence we named this protein PFR assembly factor 1, PFR-AF1.

**Figure 2:**
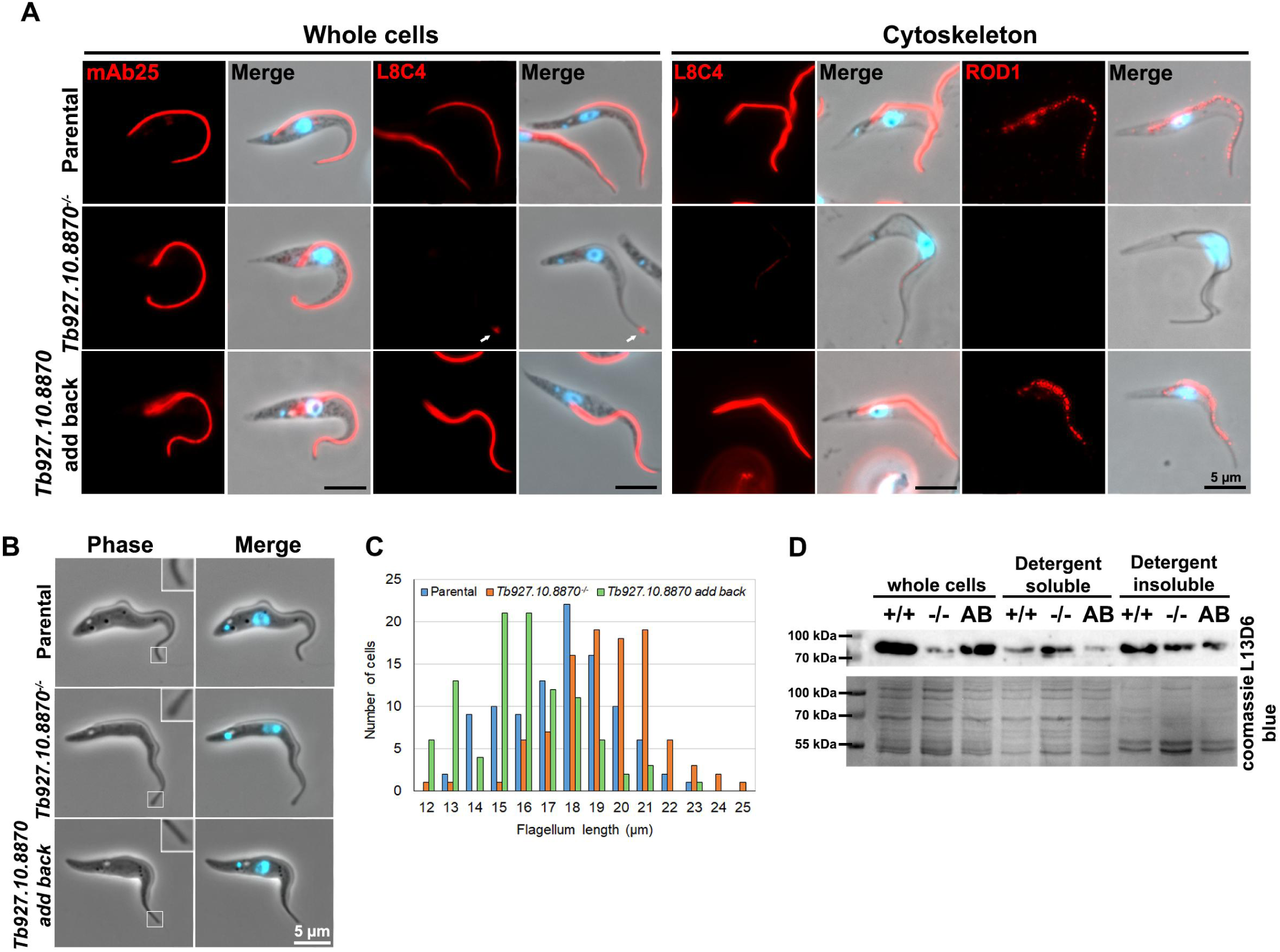
PFR assembly was disrupted in *Tb927.10.8870*^*-/-*^ mutant. (A) Immunofluorescence was performed using monoclonal antibodies mAb25 (recognizing TbSAXO) and L8C4 (recognizing PFR2) on methanol fixed cells (whole cells), and L8C4 and ROD1 (recognizing the distal domain protein Tb5.20) on cytoskeleton preparations. Antibodies labelling (red) and merge images (antibody, Hoechst 33342 and phase) of parental, *Tb927.10.8870*^*-/-*^ and *Tb927.10.8870 add back* cell lines are shown. White arrow indicates bulge of PFR material at flagellum tip. Scale bars: 5 µm. (B) Images of parental, *Tb927.10.8870*^*-/-*^ and *Tb927.10.8870 add back* 1K1N live cells. Inset showing flagellar tip. Scale bar: 5 µm. (C) Histogram of flagellum length. Flagellum length was measured in 100 parental, *Tb927.10.8870*^*-/-*^ and *Tb927.10.8870 add back* 1% NP-40 extracted cells. (D) Fractionations of parental (+/+), *Tb927.10.8870*^*-/-*^ (-/-) and *Tb927.10.8870 add back* (AB) cells using 1% NP-40. Western blot was performed on whole cells, and detergent soluble and insoluble fractions using monoclonal antibody L13D6 that recognizes PFR1 and PFR2. Coomassie blue stained gel was used as a loading control.

We further analysed the effect of PFR-AF1 deletion by generating detergent resistant cytoskeletons of parental, null mutant and add back cells, which were stained with L8C4 and ROD1, a monoclonal antibody against Tb5.20 (Woodward et al., 1994), a component of the distal domain of the PFR (Figure 2A). In the null mutant cytoskeletons L8C4 signal was observed; however, the signal was much lower and had a patchy distribution in comparison to the parental and add back cells. Conversely, ROD1 signal was not observed in the null mutant cytoskeletons, whilst a spotty distribution was seen along the flagellum in both the parental and add back cells. Together this provided further evidence that the loss of PFR-AF1 affected the stability of the PFR with Tb5.20 completely lost and PFR2 much reduced on detergent treatment. We next measured the length of the flagellum in cells with one flagellum in the parental, null mutant and add back cells (Figure 2C). The parental cells had a mean flagellum length of 18.1 µm (n=100), whereas the null mutant had a slightly longer mean flagellum length, 19.9 µm (n=100) and the add back cells had a shorter mean flagellum length, 16.4 µm (n=100). This suggests that there is a potential connection between PFR assembly and flagellum length.

Given the disruption to the PFR we investigated the subcellular distribution of PFR1 and PFR2 by using the L13D6 monoclonal antibody on western blots of whole cell, detergent soluble, and detergent insoluble lysates from the parental, null mutant and add back cells (Figure 2D). In the whole cell lysates, the amount of PFR1 and 2 detected was much lower in the null mutant that in the parental and add back cells. In addition, there was a substantial fraction of PFR1 and 2 protein detected in the soluble fraction of the null mutant, whereas in the parental and add back cells PFR1 and 2 were mainly found in the insoluble fraction. This again showed that the loss of PFR-AF1 disrupted PFR formation with a substantial amount of PFR1 and 2 not integrated into the structure.

To investigate the PFR assembly defect in more detail we examined the parental, null mutant and add back cells by thin section transmission electron microscopy (Figure 3Aa-f). In transverse sections across the flagellum of parental and add back cells we observed the expected 9+2 microtubule axoneme with the PFR alongside. The PFR was attached to microtubule doublets 4 and 7 by a linker structure and the paracrystalline nature of the PFR was observed in both longitudinal and transverse sections. In the null mutant cells, the microtubule axoneme appeared intact and similar to that observed the parental and add back cells. In *T. brucei* the central pair is aligned parallel to the PFR and this alignment was unaffected in the null mutant (Gadelha et al., 2006). However, the PFR structure was severely affected (Figure 3Ac-d). The bulge observed by light and fluorescence microscopy was clearly seen by TEM and consisted of a large amorphous collection of electron dense fibres that likely correspond to mis- or unassembled PFR components (Figure 3Ad). However, in cross sections containing a small, mis-formed PFR the linkers connecting the PFR to the microtubule doublets 4 and 7 were still present, suggesting that PFR-AF1 was not required for linker assembly into the axoneme. To quantify the changes we observed in the PFR, we defined four categories of PFR structure, i) normal, ii) proximal domain only, iii) reduced and iv) enlarged (Figure 3B). For the parental and add back cells the majority of flagellar cross sections had a normal PFR structure, whereas for the null mutant the majority of the cross sections had the proximal PFR domain only with a fraction also having a further reduced PFR or an enlarged PFR. This provides further evidence that PFR-AF1 was required for the correct formation of the PFR.

**Figure 3:**
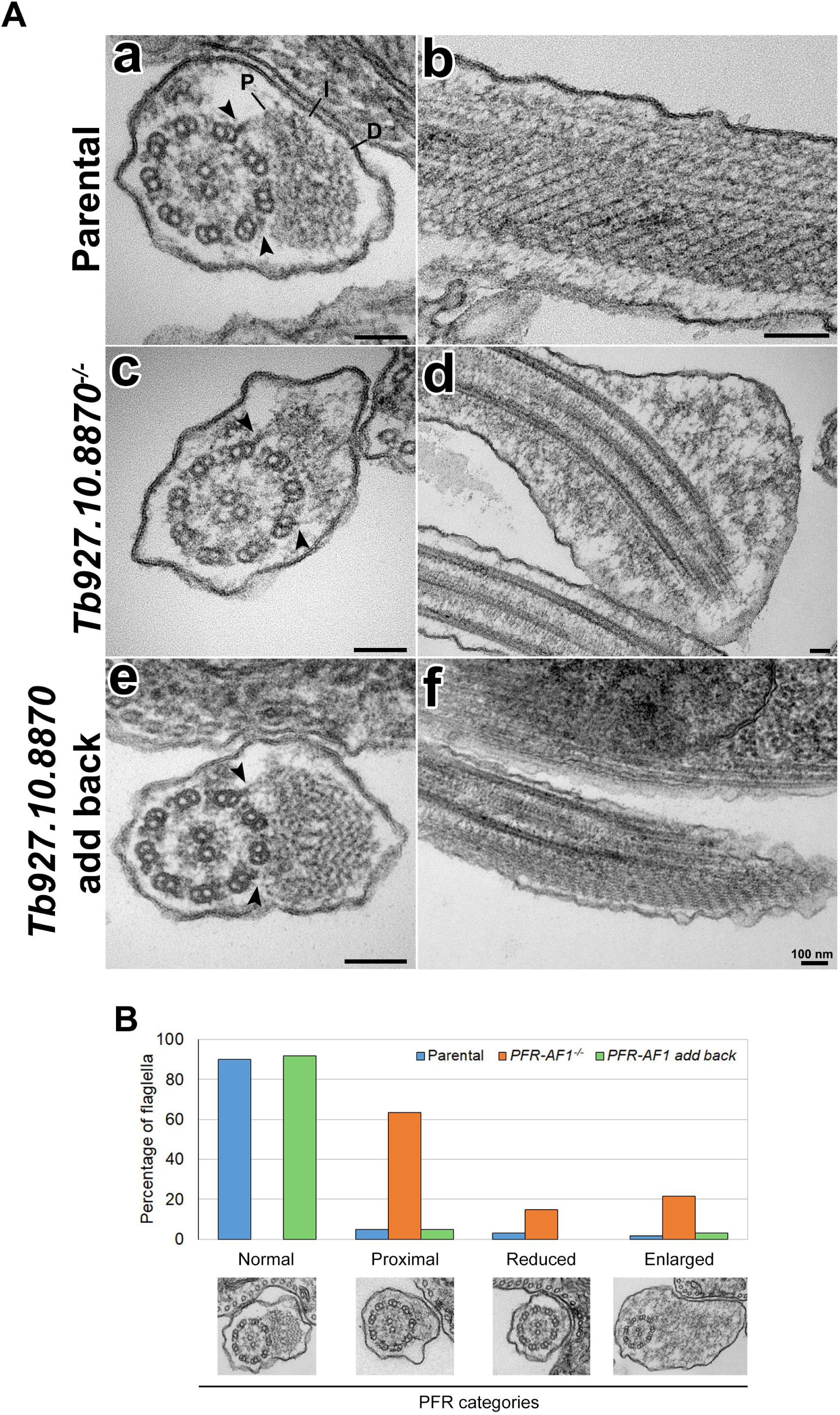
PFR structure was perturbed in *PFR-AF1*^*-/-*^ mutant. (A) TEM images of transversal (left) and longitudinal (right) flagellum sections of parental (a, b), *Tb927.10.8870*^*-/-*^ (c, d) and *Tb927.10.8870 add back* (e, f) cell lines. Arrows show the bridges that connect PFR to the axoneme. In A,a PFR domains are indicated as P (proximal), I (intermediate) and D (distal). Scale bars: 100 nm. (B) TEM observation of 60 random flagellum sections of parental, *PFR-AF1*^*-/-*^ and *PFR-AF1* add back cell lines were used to separate PFR phenotypes into four categories: normal, proximal, reduced and enlarged. Under columns of each category, a representative TEM image is shown.

### PFR-AF1 interacts with a cytoplasmic coiled coil protein

The cytoplasmic localisation of PFR-AF1 suggested that this protein is unlikely to be involved in the transport of PFR protein components into and along the flagellum or their incorporation in the flagellum. The assembly of certain axonemal components such as the outer dynein arms occur in the cytoplasm before the assembled complex is transported into the flagellum (Desai et al., 2018). To investigate whether PFR-AF1 interacts with any PFR proteins we performed immunoprecipitation using the mNG trap and mass spectrometry to identify interacting protein partners. We compared the enrichment of proteins identified in the cell line expressing PFR-AF1::mNG with that of a cell line expressing a soluble mNG (Figure 4A). PFR-AF1 was clearly enriched and another protein Tb927.7.1360 was also enriched to a similar degree. Two further proteins, Tb927.5.3060 and Tb927.11.3510 were enriched but not as significantly as Tb927.7.1360 and had low Mascot scores (<12). However, our approach did not identify any known PFR proteins.

**Figure 4:**
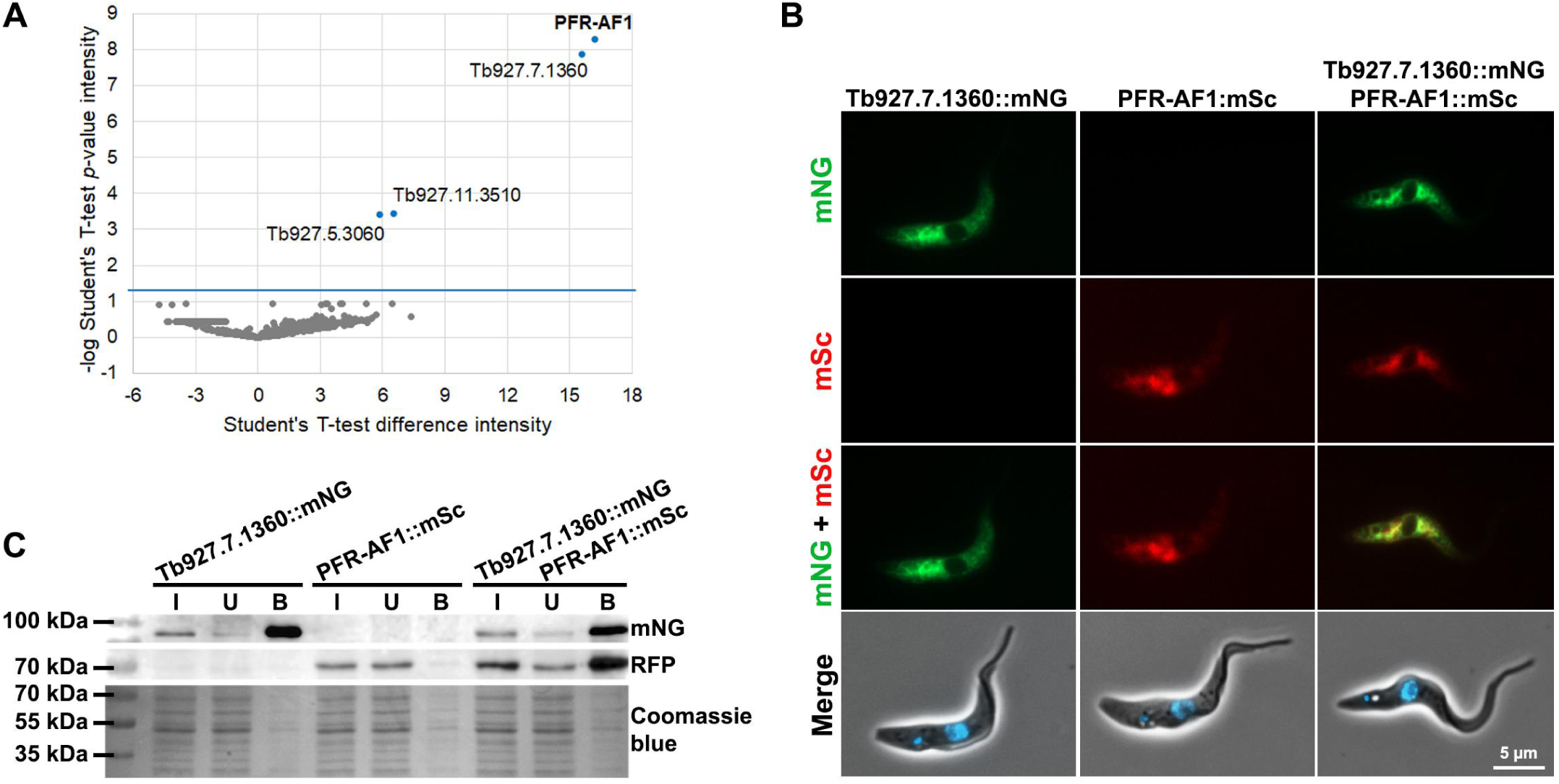
Tb927.7.1360 was a binding partner of PFR-AF1. (A) Identification of PFR-AF1 interacting proteins by mass spectrometry. Volcano plot of Student t-test difference in intensity versus p-value of intensity. (B) Fluorescence images of cell lines expressing either Tb927.7.1360::mNG (green), PFR-AF1::mSc (red) or both. Scale bars: 5 µm. (C) Immunoprecipitations with cell lines expressing Tb927.7.1360::mNG, PFR-AF1::mSc and, Tb927.7.1360::mNG and PFR-AF1::mSc with the mNG trap. Western blots were performed on the input (I), unbound (U), and bound (B) material for each cell line using anti-mNG and anti-RFP antibodies with a Coomassie blue stained gel to show total protein present. PFR-AF1::mSc was only detected when Tb927.7.1360::mNG was present.

An alternative explanation for the effect of PFR-AF1 deletion on PFR assembly is that it is a cytoplasmic factor required for the specific translation of PFR proteins and its loss results in reduced PFR protein expression. To test this hypothesis, we treated the cells with MG132, an inhibitor of the proteasome, which will result in the stabilisation of proteins that would normally be degraded but has no known effect on translation. If PFR-AF1 is important for PFR protein translation, then the addition of MG132 to the null mutant will not alter that amount of PFR protein in the cell. However, after addition of MG132 to the null mutant for 4/8 hours we saw an increase in the amount of PFR1 and 2 (Figure S2B), suggesting that PFR 1 and 2 were being degraded in the PFR-AF1 null mutant by the proteasome and that PFR-AF1 did not have a role in PFR protein translation but instead was important for PFR protein stability.

### Tb927.7.1360 is also required for PFR assembly

Given the significant enrichment of Tb927.7.1360 and that its localisation in the TrypTag database was similar to that of PFR-AF1 we decided to focus on this protein. Tb927.7.1360 was predicted to form a single coiled coil domain and was restricted to the kinetoplastids and not found in other organisms. To confirm that PFR-AF1 and Tb927.7.1360 interacted we performed a co-immunoprecipitation using Tb927.7.1360 as the bait. We generated three cell lines, one expressing Tb927.7.1360 endogenously tagged with mNG, another cell line expressing PFR-AF1 tagged with mScarlet and finally one expressing both Tb927.7.1360::mNG and PFR-AF1::mSc, both the tagged proteins had the expected cytoplasmic localisation (Figure 4B). On immunoprecipitation using the mNG trap with these three cell lines PFR-AF1::mSc was only detected when Tb927.7.1360::mNG was present, indicating that these two proteins specifically interact (Figure 4C).

To understand the function of Tb927.7.1360 we generated a cell line in which both alleles of Tb927.7.1360 were replaced with antibiotic resistance markers. Loss of the Tb927.7.1360 open reading frame and integration of the resistance makers was confirmed by PCR (Figure S3). The growth rate of the Tb927.7.1360 null mutant was compared to that of the parental cell line and we found that it was consistently slower growing (Figure 5A). Imaging of the Tb927.7.1360 null mutant stained with the DNA stain, Hoechst 33342 showed a similar range of cell cycle defects observed with the deletion of PFR-AF1 (Figure 5B). In addition, we noticed that there was a bulge at the tip of the flagellum of ∼52% (n=240) of the one flagellum cells in the Tb927.7.1360 null mutant, again as previously seen with loss of PFR-AF1. Together this suggested that Tb927.7.1360 was likely affecting PFR formation.

**Figure 5:**
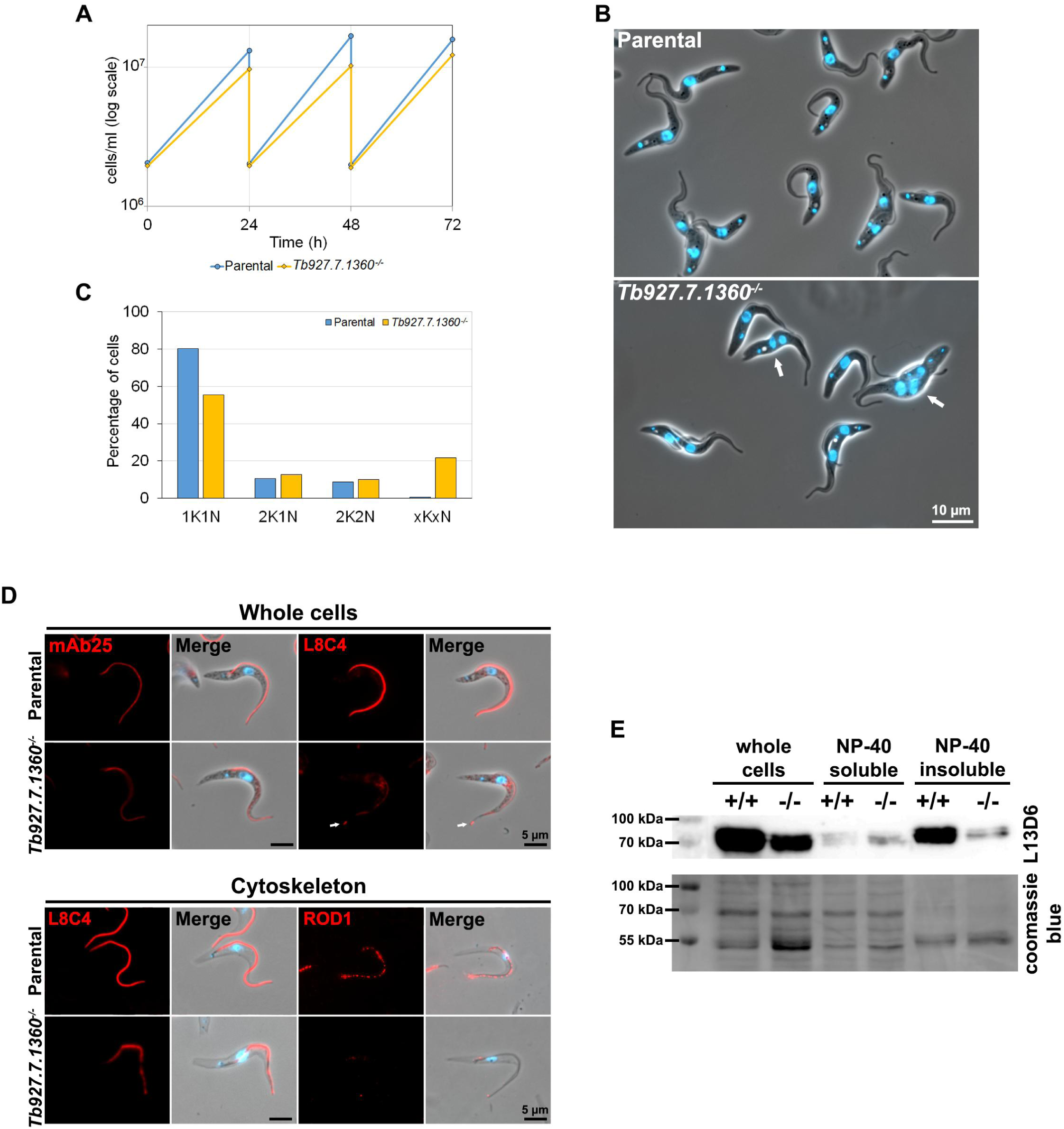
Tb927.7.1360 deletion disrupted PFR formation. (A) Growth curves of parental and *Tb927.7.1360*^*-/-*^ cell lines over 72 hours. (B) Merge of phase and Hoechst images of parental and *Tb927.7.1360*^*-/-*^ live cells. White arrows indicate abnormal cells. Scale bar: 10 µm. (C) Nuclear (N) and kinetoplast (K) DNA was counted in 200 cells using Hoechst 33342 stained fluorescence images of parental and *Tb927.7.1360*^*-/-*^ cell lines. All cells with abnormal numbers of K and N were counted as “xKxN”. (D) Immunofluorescence was performed using monoclonal antibodies mAb25 (recognizing TbSAXO) and L8C4 (recognizing PFR2) on methanol fixed cells (whole cells), and L8C4 and ROD1 (recognizing the distal domain protein Tb5.20) on cytoskeleton preparations. Antibodies labelling (red) and merge images (antibody, Hoechst 33342 and phase) of parental and *Tb927.7.1360*^*-/-*^ cell lines are shown. Scale bars: 5 µm. (E) Fractionations of parental (+/+) and *Tb927.7.1360*^*-/-*^ (-/-) cells using 1% NP-40. Western blot was performed on whole cells, and detergent soluble and insoluble fractions using monoclonal antibody L13D6 that recognizes PFR1 and PFR2. Coomassie blue stained gel was used as a loading control.

We investigated the effect of Tb927.7.1360 deletion on the structure of the axoneme and PFR by staining the cells with the axoneme antibody mAb25 and the PFR antibody L8C4 (Figure 5D). There was no difference in staining pattern of mAb25 between the parental and Tb927.7.1360 null mutant cells as found with the deletion of PFR-AF1. In the parental cells L8C4 strongly stained the PFR within the flagellum. However, in the null mutant the L8C4 signal in the flagellum was no longer evenly distributed with patches of stronger staining interspersed with regions of very weak staining. We investigated the defect in PFR formation further by staining cytoskeletons with L8C4 and ROD1 (Figure 5D). In parental cytoskeletons L8C4 gave a strong signal along the flagellum, whereas in the null mutant the signal was weaker with distinct gaps. ROD1 as expected gave a patchy distribution along the flagellum in parental cytoskeletons; however, in the null mutant cytoskeletons ROD1 was barely detectable with only a spot of signal at the proximal and distal end of the flagellum seen. Given the effect Tb927.7.1360 had on PFR assembly, we named this protein PFR assembly factor 2, PFR-AF2. To complement the microscopy, we used detergent to fractionate the cell into soluble and insoluble fractions, which were analysed by western blotting with L13D6 (Figure 5E). Deletion of PFR-AF2 reduced the amount of PFR1 and 2 in the cells with a marked reduction seen in the insoluble fraction. In the null mutant soluble fraction there was more PFR1 and PFR2 present than in the equivalent parental fraction. The similar phenotypes observed on deletion of PFR-AF1 and PFR-AF2 and that they interact suggest that these proteins work together to enable PFR formation.

## Discussion

Extra-axonemal structures are found in many eukaryotic flagella including the outer dense fibres and fibrous sheath of sperm and the mastigonemes of *Chlamydomonas*. Initial research on these structures described their structure and assembly kinetics, followed by cataloguing and functionally analysing their constituent parts (Eddy et al., 2003; Irons and Clermont, 1982a; Irons and Clermont, 1982b; Nakamura et al., 1996; Ringo, 1967; Witman et al., 1972); however, little work has focussed on the mechanism of assembly of these structures. Due to the lack of ribosomes in the flagellum all the proteins required for flagellar assembly must undergo a multi-step process before being finally incorporated into the flagellum, including translation in the cytoplasm, assembly and post-translational modification if required, sorting and delivery to the flagellum, transport along the flagellum, and assembly into the flagellum structure. These processes occur across three main compartments, the cytoplasm, the transition fibre-transition zone, and the flagellum. The assembly of extra-axonemal structures including the PFR and the outer dense fibres is likely to follow a similar pattern (Bastin et al., 1999b).

Here, we have shown that two proteins, PFR-AF1 and PFR-AF2, form a cytoplasmic complex specifically important for PFR assembly. These are the first cytoplasmic proteins to be associated with PFR formation. Deletion of either of these proteins had similar phenotypes, the null mutants had a reduced amount of PFR1 and PFR2, were unable to assemble an intact PFR, grew slower and abnormal cells were observed. To date the only other proteins important for PFR formation are components of either the PFR itself (PFR1, PFR2, calmodulin), or axoneme and basal body (Kif9b), or the transition fibres (FOPL) (Bastin et al., 1998; Bastin et al., 1999a; Demonchy et al., 2009; Ginger et al., 2013; Harmer et al., 2018; Maga et al., 1999; Santrich et al., 1997). The deletion of both PFR-AF1 and PFR-AF2 resulted in the accumulation of PFR proteins at the tip of the flagellum, showing that PFR proteins were still able to be trafficked into the flagellum and transported to the flagellum tip, but once there they were not effectively assembled into the PFR structure. This suggests that the action of PFR-AF1 and PFR-AF2 is required for PFR proteins to be competent for PFR formation or that without PFR-AF1 or PFR-AF2 not all PFR components were able to reach the flagellum so disrupting PFR formation.

The residual PFR structure observed by TEM when PFR-AF1 was deleted was similar to that observed with PFR2 RNAi knockdown, yet PFR2 was still present in the PFR of the flagellum in the null mutants so it is likely that composition of this residual structure is different to that observed in the PFR2 RNAi mutant (Bastin et al., 1998). Interestingly, in TEM cross sections of the flagellum in the Kif9b, calmodulin and FOPL RNAi mutants there were often multiple, individually discernible PFR cross sections that though disordered were similar in outline and size to a normal PFR cross section, suggesting that the ability to assemble a PFR type structure was present and that the failure was a lack of integration of these structures into a single continuous PFR alongside the axoneme (Demonchy et al., 2009; Ginger et al., 2013; Harmer et al., 2018). However, in the PFR-AF1/2 deletion mutants, a normal sized PFR structure was never observed, suggesting that the cytoplasmic PFR-AF1 and PFR-AF2 act upstream of Kif9b, calmodulin and FOPL in the PFR formation process. This fits well with the concept of flagellum assembly being composed of multiple processes across different locations and suggests that FOPL, which localises to transitional fibres on the basal body, may act to regulate entry of PFR proteins into the flagellum with KIF9B/calmodulin required for their assembly within the flagellum.

In sperm, the ubiquitin conjugating enzyme, UBE2B was shown to be important for the positioning and assembly of the longitudinal columns of the fibrous sheath (Escalier, 2003). Addition of ubiquitin can have a range of effects on a protein, including targeting it for degradation or altering its location in the cell; however, its specific function in fibrous sheath assembly is unknown. The TrypTag project has identified two ubiquitin associated proteins that localise to the axoneme and others that localise to the flagellar cytoplasm but none specific for the PFR (Dean et al., 2017). In addition, no further proteins have been identified to date that are important for assembly of the sperm extra-axonemal structures beyond those that are integral components of them. However, interrogating the function of proteins involved in outer dense fibre/fibrous sheath assembly in sperm is difficult as these are terminally differentiated cells and therefore the PFR and its assembly mechanism provides a paradigm that will be useful when considering extra-axonemal structure assembly in other organisms.

The PFR has a regular, ordered structure and it is unlikely that this structure would arise spontaneously, suggesting additional proteins are required to assist its assembly. Based on the assembly model outlined above one can conjecture a number of different functions for these additional proteins including cytoplasmic assembly, post translational modification, transport, and flagellar based chaperones. As the PFR contains nearly 200 proteins (Dean et al., 2017), it would be unlikely that each would have its own set of specific associated factors; therefore, PFR-AF1 and PFR-AF2 would be more likely employed assisting the assembly/modification/transport of multi-subunit complexes that are then incorporated into the PFR. This has clear parallels with dynein arm assembly. The assembly mechanism for the dynein arms that drive and coordinate flagellum movement has been well studied. This process has a cytoplasmic step during which the dynein arm proteins are folded and assembled into a complex before undergoing further maturation and delivery to the flagellum (Desai et al., 2018). This cytoplasmic step requires an expanding list of dynein arm assembly factors (currently 11 – DNAAF1/LRRC50, DNAAF2/Kintoun, DNAAF3, MOT48, HEATR2, LRRC6, DYX1C1/DNAAF4, PIH1D3/Twister, SPAG1, ZMYND10 (Cho et al., 2018; Desai et al., 2018; Horani et al., 2012; Loges et al., 2009; Mitchison et al., 2012; Mitchison et al., 2012; Olcese et al., 2017; Omran et al., 2008; Tarkar et al., 2013)) that act as chaperones to ensure successful folding and assembly of the dynein arm complex. In the PFR-AF1 null mutant PFR1 and PFR2 protein amount increased when the cells were treated with the proteasome inhibitor MG132, suggesting that without PFR-AF1 PFR1/PFR2 were unstable and degraded. This phenomenon has also observed for dynein heavy chains when DNAAFs were depleted such as DNAAF3 (Mitchison et al., 2012).

The role of the PFR-AF1/PFR-AF2 complex in PFR formation still needs to be defined. The cytoplasmic localisation of these proteins and the possible roles for PFR-AFs discussed above may indicate that PFR-AF1/PFR-AF2 are involved in the assembly of multi-subunit complexes. For example, do PFR-AF1/PFR-AF2 act as a scaffold onto which PFR proteins can assemble before disengaging to allow the PFR unit to be transported to the flagellum? However, an enrichment of PFR proteins in the PFR-AF1 immunoprecipitation was not seen. Any interaction between the PFR-AF1/PFR-AF2 complex and any PFR proteins is likely to be transient and the methods used here were not sensitive enough to detect any such interaction. Moreover, we do not know which of the nearly 200 PFR proteins are present in these putative PFR protein complexes, there is evidence for interaction between PFR proteins; however, none of these interactions involved PFR1 and PFR2 (Lacomble et al., 2009; Portman and Gull, 2010). There is also the potential for multiple types of unit as the composition of the proximal domain and mid/distal domain are different with the former able to form without PFR2.

PFR-AF1 and PFR-AF2, 34 kDa and 42 kDa respectively, are predicted to form coiled coils. PFR-AF2 is restricted to the kinetoplastids, whereas PFR-AF1 was identified as containing a taxilin PFAM domain and this domain is implicated in vesicular trafficking by binding to syntaxins. However, the cytoplasmic localisation of PFR-AF1 does not support a role in vesicular trafficking and this domain may therefore have evolved to bind a different set of partners in the kinetoplastids. The PFR is specific to *Euglenozoa* but it can vary in appearance greatly between different species with *Angomonas deanei* and *Strigomonas culicis* having a much-reduced PFR structure that does not contain PFR2 (Gadelha et al., 2005; Motta et al., 2013). However, both PFR-AF1 and PFR-AF2 are conserved in these species and it is likely that even with a reduced PFR these proteins are required for its correct formation.

In summary, our data fit with the concept of multiple flagellum assembly processes occurring across different cellular compartments. The discovery of analogous set of positioned processes for extra-axonemal structures suggests that this is common mechanism to deal with complications of building a complex structure whose site of assembly can be many microns from the cytoplasm.

## Materials and Methods

### Cell culture

*Trypanosoma brucei* TREU927 procyclic forms containing the plasmid pJ1339 that expresses T7 RNA polymerase, tet repressor and Cas9 were grown at 28°C in SDM-79 medium supplemented with 10% FCS (Brun and Schönenberger, 1979). Cell concentration was determined in a Z2 Coulter Counter particle counter.

### Generation of deletion constructs, tagging constructs and add back constructs

Constructs for endogenous gene tagging and gene deletion mediated by CRISPR-Cas9 were generated as described by (Beneke et al., 2017). To generate the *Tb927.10.8870* add back construct the *Tb927.10.8870* open reading frame was cloned into plasmid pJ1313 using the SpeI and BamHI restriction sites, which resulted in a fusion protein with an N-terminal Ty tag. pJ1313 is a modified version of p3605 (de Freitas Nascimento et al., 2018) in which additional SpeI sites have been removed and the 3Ty::GS::mNG::GS::3Ty cassette has been cloned into the HindIII/BamHI sites. This plasmid will support constitutive gene expression and enables the generation of proteins tagged with 3Ty and/or mNG at either the N- or the C-terminus. All constructs were electroporated using Nucleofector 2b device and Program X-001 (Dean et al., 2015).

### Electron microscopy

#### TEM

Cells were fixed by addition of glutaraldehyde to the final concentration of 2.5% into the cultures. Cells were harvested by centrifugation at 800xg for 5 minutes and primary fixative (2.5% glutaraldehyde, 2% formaldehyde, 0.1 M sodium phosphate buffer pH 7.0) was added without disturbing the pellet. After 1 hour at room temperature, samples were washed three times in 0.1 M sodium phosphate buffer pH 7 for five minutes and post-fixed in 1% osmium tetroxide in 0.1 M sodium phosphate buffer pH 7 for 90 minutes at room temperature under agitation. Samples were washed in distilled water three times for 5 minutes and stained in 2% uranyl acetate for 12 hours at 4°C. Samples were washed three times in water and dehydrated in increasing concentrations of acetone (30%, 50%, 70%, 90% v/v in distilled water, followed by three times in 100% acetone) and embedded in Agar-100 resin. Thin sections were post-stained in 2% uranyl acetate and 3% lead citrate. Images were obtained using Hitachi H-7650, FEI Tecnai G2 Spirit or Jeol 1400Flash, operated at 120 kV.

#### SEM

Fixation was performed by adding glutaraldehyde to final concentration of 2.5% into the cell culture. Cells were harvested by centrifugation at 800xg for 5 minutes and primary fixative was added (2.5% glutaraldehyde in PBS). After two hours, cells were washed twice in PBS and settled onto round glass coverslips for 5 minutes. Coverslips were washed two times in PBS and samples dehydration was performed using increasing concentrations of ethanol (30%, 50%, 70% and 90% v/v in distilled water, followed by three times in 100% ethanol) for five minutes in each step. Samples were dried using a critical point dryer. Coverslips were mounted onto SEM stubs using silver DAG and coated with gold using a sputter coater. Images were acquired on a Hitachi S-3400N scanning electron microscope.

#### Antibodies

The source of antibodies used in this work and their dilutions were as follow. Monoclonal mNeonGreen antibody from Chromotek (32F6) diluted 1:100 for Western blotting. Monoclonal RFP antibody from Chromotek (6G6) diluted 1:1000 for Western blotting. Monoclonal BB2 (Bastin et al., 1996) diluted 1:1000 for Western blotting or 1:100 for immunofluorescence. Monoclonal L8C4 (Kohl et al., 1999) diluted 1:1000 for Western blotting or 1:200 for immunofluorescence. Monoclonal L13D6 (Kohl et al., 1999) diluted 1:100 for Western blotting. Monoclonal mAb25 (Dacheux et al., 2012) diluted 1:100 for immunofluorescence. Monoclonal ROD1 (Woods et al., 1989) diluted 1:200 for immunofluorescence. Secondary IgG anti-mouse conjugated to Alexa Fluor 546 from Invitrogen (A11030) diluted 1:200 for immunofluorescence. Secondary IgM anti-mouse conjugated to Alexa Fluor 546 from Invitrogen (A21045) diluted 1:250 for immunofluorescence. Secondary IgG antimouse conjugated to peroxidase from Jackson ImmunoResearch (715-035-150) diluted 1:1000 for Western blotting.

#### Cell fractionation

2×10^6^ cells were harvested by centrifugation at 800xg for 5 minutes, washed in PBS and incubated with 50 µl of 1% NP-40 in PEME (0.1 M PIPES pH 6.9, 2 mM EGTA, 1 mM MgSO4, 0.1 mM EDTA) containing protease inhibitors for 5 minutes on ice. To obtain whole cells samples, after the treatment, 50 µl of 2x Laemmli buffer was added to cell lysate. Soluble and insoluble fractions were separated by centrifugation at 20000xg for 10 minutes at 4°C. To the soluble fraction, 50 µl of 2x Laemmli buffer was added. The insoluble fraction was resuspended into 50 µl of 1% NP-40 in PEME and 50 µl of 2x Laemmli buffer. All samples were incubated at 100°C for 5 minutes.

#### Western blotting

10^6^ cell equivalents were loaded onto 12% SDS-PAGE gel. After transfer, membranes were blocked with blocking solution (3% milk, 0.05% Tween 20 in PBS) for 1 hour, incubated with primary antibody diluted in blocking solution for 1 hour, washed three times in blocking solution and incubated with secondary antibody diluted in blocking solution for 1 hour. Proteins were detected by ECL.

#### Immunofluorescence microscopy

Cells were harvested by centrifugation at 800xg for 5 minutes, washed in vPBS and settled onto glass slides for 5 minutes. For whole cells preparations, cells were fixed in −20°C methanol for 20 minutes and rehydrated in PBS for 10 minutes. For cytoskeleton preparations, cells were first treated with 1% NP-40 in PEME for 5 minutes, washed in PBS and fixed in −20°C methanol for 20 minutes. After fixation, the slides were blocked for 1 hour at room temperature with blocking solution (1% BSA in PBS) and incubated in primary antibody diluted in blocking solution for 1 hour. After washing three times in PBS, slides were incubated in secondary antibody diluted in blocking solution for 1 hour. After three washes in PBS, slides were incubated in 20 μg/ml Hoechst 33342 (Sigma-Aldrich) in PBS, washed in PBS and mounted before imaging. Images were taken using a Zeiss Axio Imager.Z1 microscope equipped with an ORCA-Flash 4.0 CMOS camera using a Plan-Apochromat 100x/1.4 NA oil lens. Images were acquired and analysed with ZEN 2 PRO and assembled for publication in Adobe Photoshop CS6.

#### Immunoprecipitation

2×10^8^ cells were harvested by centrifugation at 800xg for 5 minutes, washed in PBS and resuspended into 1 ml of lysis buffer (0.2% NP-40, 10 mM Tris-HCl pH 7.5, 150 mM NaCl) for 5 minutes on ice. Soluble proteins were separated by centrifugation at 16000xg for 10 minutes at 4°C. 25 µl of mNeonGreen-trap magnetic agarose previously washed in washing buffer (10 mM Tris-HCl pH 7.5, 150 mM NaCl) were incubated with 500 µl soluble protein lysate (final concentration of 10^8^ cells) for 30 minutes at 4°C tumbling end-over-end. Beads were magnetically separated and washed twice with washing buffer. Samples were eluted into 100 µl 1x Laemmli buffer for Western blotting analysis or into 50 µL of 4M urea for mass spectrometry analysis.

#### Mass spectrometry

Eluates from the immunoprecipitation were reduced, alkylated and then digested overnight with trypsin. The peptides were separated on an Ultimate 3000 UHPLC system and then directly electrosprayed into the coupled QExactive mass spectrometer. The raw data was acquired on the mass spectrometer in a data-dependent mode and full scan MS spectra were acquired in the Orbitrap. The raw data files were processed using MaxQuant, integrated with the Andromeda search engine. For protein identification, peak lists were searched against *T. brucei* database and a list of common contaminants. Protein and PSM false discovery rate were set at 0.01. Protein group txt file was them imported to Perseus 1.5.2.4, to perform student t-test.

## Acknowledgements

We would like to thank Dr Artur de Castro Neto (Oxford Brookes University) for his technical assistance. We are grateful for Prof Derrick Robinson (University of Bordeaux) for providing the mAb25 antibody. We would also like to thank Svenja Hester (University of Oxford) for assistance with the proteomics and Dr Flavia Moreira-Leite (Oxford Brookes University) and the Oxford Brookes Bioimaging Unit for assistance with the electron microscopy. This work was initiated in the laboratory of Prof Keith Gull (University of Oxford) and we are grateful for his advice and input throughout this project.

## Competing interests

No competing interests declared.

## Funding

This work was funded by MRC Newton Fund award (MR/N017323/1) to SV and NCS, and the Wellcome Trust (104627/Z/14/Z, 108445/Z/15/Z) to Prof Keith Gull. FAPERJ and CNPq support the Cell Structure Laboratory in Brazil (NCS and WS). MJRB was supported by an Oxford Brookes University Research Fellowship.

---

Figure S1. (A) Schematic and PCR confirmation of *Tb927.10.8870* gene deletion. gDNA from *Tb927.10.8870*^*-/-*^ mutant and the parental cells was analysed by PCR. PCR confirmed that *Tb927.10.8870* ORF was no longer present in the null mutant and that the resistance markers had integrated correctly. (B) Confirmation of Ty::Tb927.10.8870 expression in add back cell line. Western blot was performed on whole cell lysates of parental, *Tb927.10.8870*^*-/-*^ and *Tb927.10.8870 add back* cell lines using monoclonal antibody BB2 that recognises the Ty tag. Coomassie stained gel was used as a loading control. (C) Confirmation of Ty::Tb927.10.8870 localisation to the cytoplasm in the add back cell line. Immunofluorescence was performed using the BB2 antibody on methanol fixed whole cells. Phase, BB2 labelling (red) and merge images (antibody, Hoechst 33342 and phase) of parental and *Tb927.10.8870*^*-/-*^ and *Tb927.10.8870 add back* cell lines are shown.

Figure S2. (A) SEM images of parental and *Tb927.10.8870*^*-/-*^ cells. The bulge is clearly present at the tip of the flagellum White arrows indicate the bulge at the tip of the flagellum. (B) Proteasome inhibitor MG132 was added to parental, *Tb927.10.8870*^*-/-*^ and *Tb927.10.8870 add back* cell lines for 8 *Tb927.10.8870 add back* cell lines using monoclonal antibody L13D6 that recognizes PFR1 and PFR2. Coomassie blue stained gel was used as a loading control. Addition of MG132 caused an increase in PFR1 and PFR2 amount in *Tb927.10.8870*^*-/-*^ mutant.

Figure S3. (A) Schematic and PCR confirmation of *Tb927.7.1360* gene deletion. gDNA from 4 *Tb927.7.1360*^*-/-*^ mutant clones and the parental cells was analysed by PCR. PCR confirmed that *Tb927.7.1360* ORF was no longer present in the null mutant clones and that the resistance markers had integrated correctly. *Tb927.7.1360*^*-/-*^ clone 1 was used for all subsequent experiments.

