## Supplementary figures and images for "Control of assembly of extra-axonemal structures: the paraflagellar rod of trypanosomes"

### Supplementary Figures

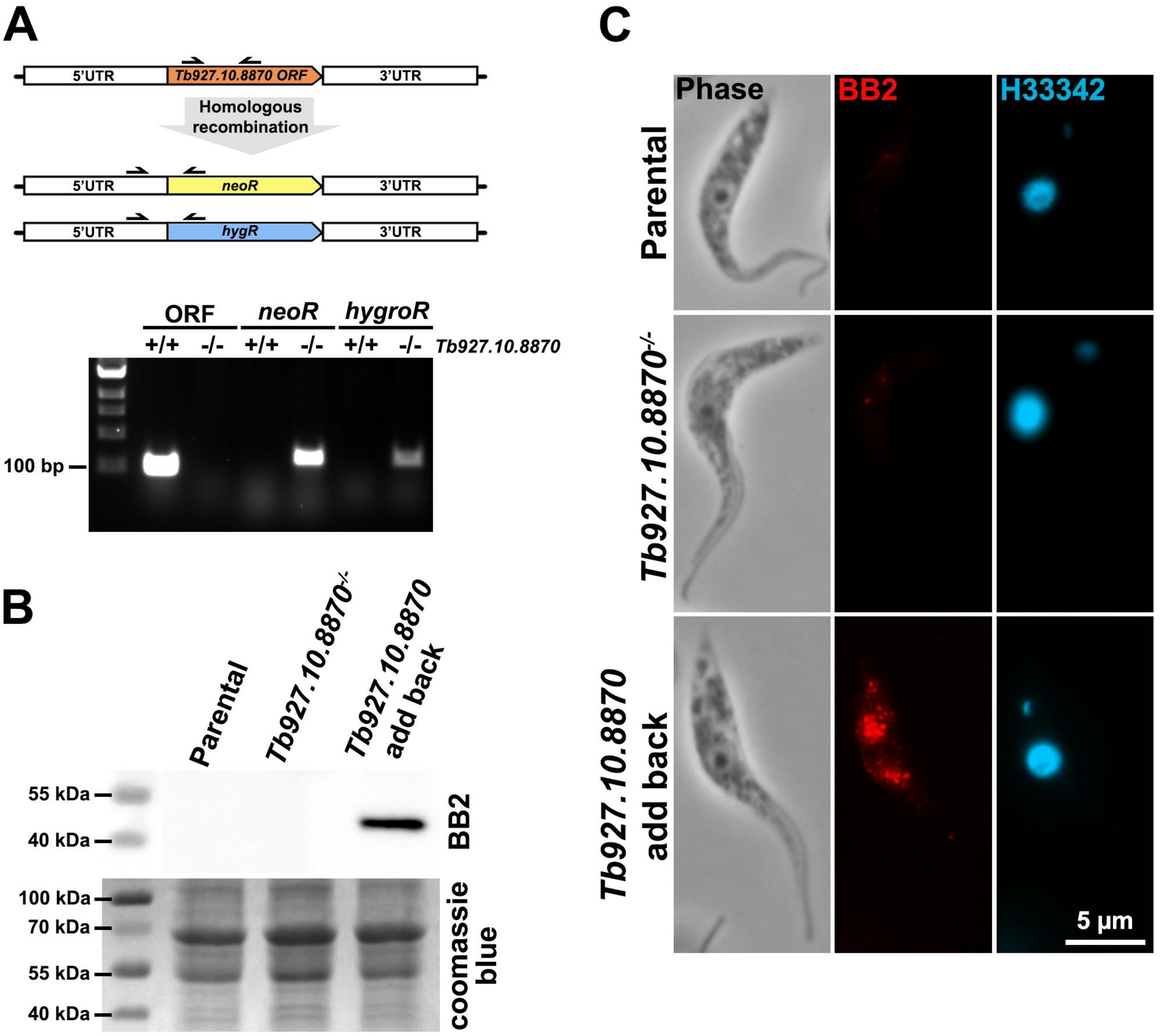

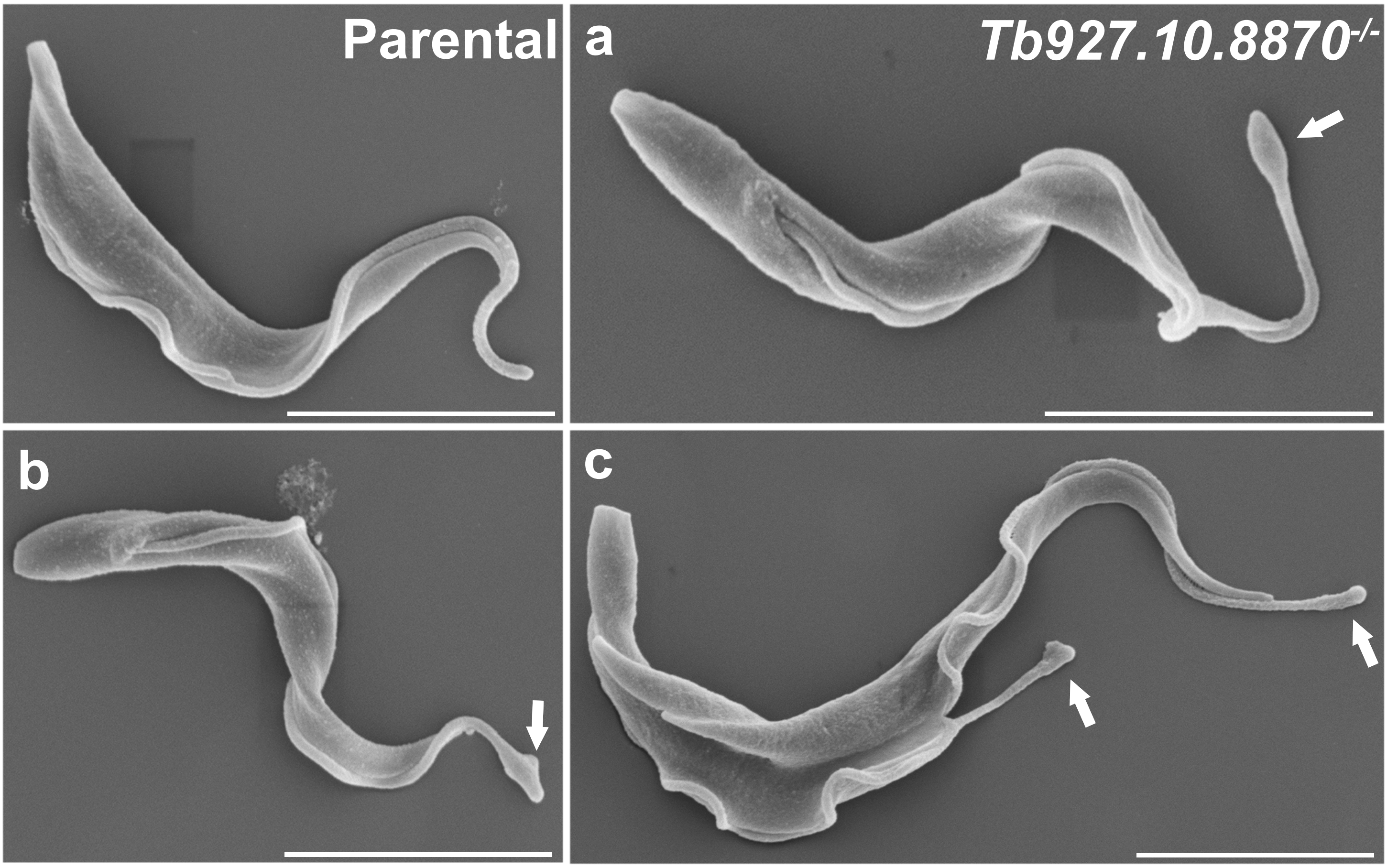

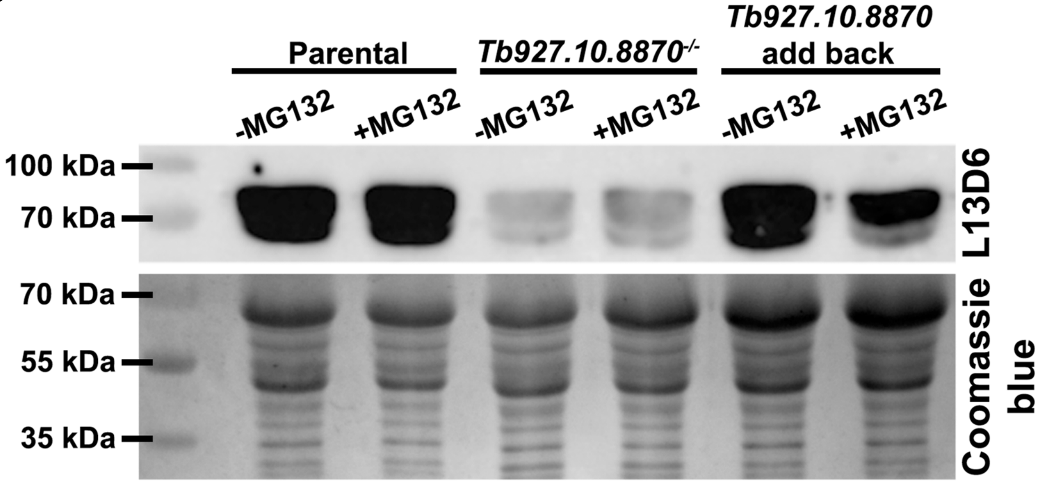

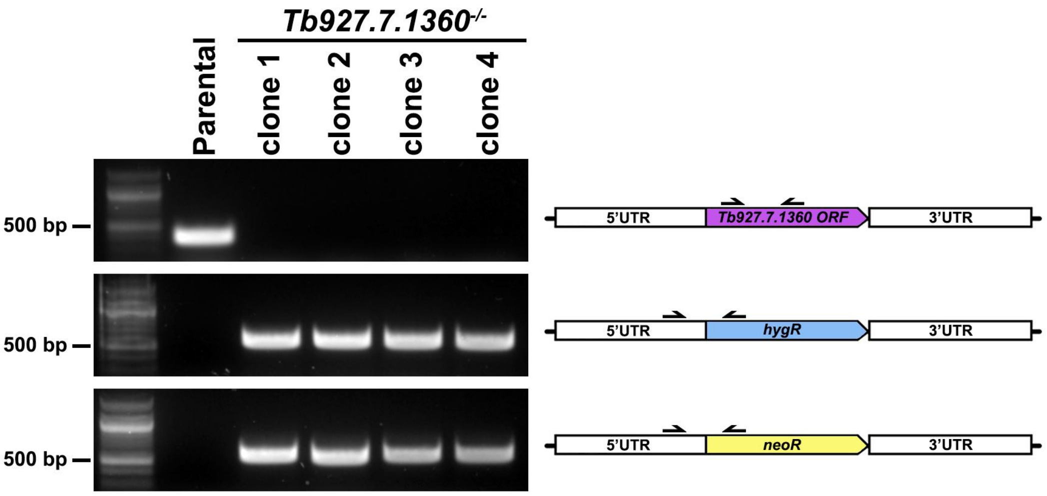
